# Early Life Exposure to Queen Mandibular Pheromone Mediates Persistent Transcriptional Changes in the Brain of Honey bee Foragers

**DOI:** 10.1101/2022.07.07.499229

**Authors:** Tianfei Peng, Anissa Kennedy, Yongqiang Wu, Susanne Foitzik, Christoph Grüter

**Affiliations:** Institute of Molecular and Organismic Evolution, Johannes Gutenberg University of Mainz, Biozentrum I, Hanns Dieter Hüsch Weg 15, 55128 Mainz, Germany; School of Biological Sciences, University of Bristol, 24 Tyndall Avenue BS8 1TQ Bristol, UK

**Keywords:** sensitive period, maturation, division of labor, signaling

## Abstract

How behavior in insect societies is regulated remains a fundamental question in sociobiology. In hymenopteran societies, the queen plays a crucial role in regulating group behavior by affecting individual behavior, physiology, and lifespan through worker gene expression. Honey bee (*Apis mellifera*) queens signal their presence via the queen mandibular pheromone (QMP). While QMP has been shown to influence the behavior of young workers, we know little about its long-term molecular impacts on workers and whether these pheromone effects depend on an early sensitive period in the life of a worker. Here we demonstrate that QMP treatment strongly impacts long-term forager gene expression in the mushroom bodies, antennal lobes, and antennae, but only if bees were treated early in life (1-2 days of age). Contrary to our expectation, foraging activity was not impacted by QMP treatment in the long-term, but genes important for division of labor, learning, chemosensory perception and aging were differentially expressed in the antennae and brain tissues, suggesting that QMP influences diverse physiological and behavioral processes in workers. Overall, our study suggests a sensitive period early in the life of workers, where the queens’ presence or absence has strong and potentially livelong effects on transcriptional activity in the central and peripheral nervous system.

**Significance statement:** Despite our increasing understanding of how social cues affect gene expression and behavior in social animals, we still know little about the importance of sensitive periods in cue perception for the long-term regulation of gene expression. Honey bees live in complex societies and queen pheromones play a central role in the regulation of worker behavior and division of labor. We tested the exposure to queen pheromone presence and found that there is a sensitive period in the early adult life of workers. Understanding and identifying sensitive periods and their effect on long-term gene transcription in workers in response to changes in the environment will lay an important foundation for a better understanding of how queens shape colony life.

## Introduction

The ecological success of social insects depends, among other factors, on an effective division of labor and communication (1-4). Ants, social bees, and social wasps show a reproductive division of labor between queens and workers, where queens are responsible for reproduction and all other colony tasks are performed by workers of different ages. In many species, young workers focus on tasks inside the colony, such as feeding the brood, whereas older workers perform tasks outside their nest, such as foraging (5-10). The behavioral maturation from in-nest work to foraging is affected by genetic factors such as parent-specific gene expression from mothers (matrigenes) and fathers (patrigenes) (11-14) as well as sensory input from social and environmental stimuli (15-17).

The queen influences worker behavior and colony dynamics through queen pheromones, with effects ranging from attracting workers to the queen (18), suppressing reproduction in workers (19, 20) and other queens (21, 22), to altering the learning capacity of workers (23). Experiments with synthetic queen pheromones and mated versus non-mated queens have identified several bioactive components that promote altruistic behaviors and regulate caste differentiation (19, 24-27). Queen pheromones of different social insect species are often chemically similar to cuticular hydrocarbons (CHCs), suggesting they evolved from conserved signals of a common ancestor (19). Probably as a result of this, cross-exposure between queen pheromones and workers of different species results in similar behavioral responses even in non-social insects (28-30).

Honey bees represent an unusual case in that the primary queen signal is a 5-compound mixture that is secreted from the mandibular gland of queens, the so-called queen mandibular pheromone (QMP). In addition, CHCs might also act as queen pheromones in honey bees (31). QMP has both short-term and long-term effects. Short-term effects include the promotion of retinue behavior, swarm clustering, and drone attraction during mating (32), while suppressing aversive learning in young bees (23). Long-term effects of QMP comprise the inhibition of queen rearing, worker reproductive development (33, 34), lowering of the sucrose response threshold (35) and the modulation of worker activities such as the transition from nursing to foraging (32, 36). While there is ample experimental evidence to support the behavioral changes induced by QMP exposure, there is limited experimental evidence to show the upstream physiological changes that result in the subsequent behavioral changes induced by QMP. Understanding how QMP exposure influences worker neurophysiology, and how long these effects last, will provide better insights into the changes in areas related to learning and memory, the mushroom bodies (37-39) and olfaction, the antennal lobes (40), which are strongly associated with behavioral transitions.

The effects of QMP have been found to be most pronounced in young bees, which exhibit an increased sensitivity to QMP (41) and in which the expression of the biogenic amine dopamine is strongly altered shortly after QMP treatments (42, 43). However, it remains unknown whether QMP influences transcriptional activity in the long-term, *i*.*e*. when bees reach forager age, and whether the effects of queen pheromone on gene expression depends on the life stage at which an individual worker experiences it.

It is common for honey bee colonies to experience temporal queen-absence due to swarming or queen-replacement, and long-term effects of the lapse in QMP exposure on the behavior and physiology of different age-cohorts are expected, possibly depending on an early sensitive period to QMP. Sensitive periods of exposure to social maternal factors have been studied in the social and biological sciences to understand time periods or life stages in which experience shapes a gene expression or behavior to a larger extend than if experienced in other time periods or life stages (44, 45). The responses are usually most pronounced during early life exposure and can carry over to influence phenotypes such as behavior, physiology, and morphology through the adult life. Current models examine how mechanisms of behavioral plasticity allow animals to optimally respond to experience across the lifespan (46-48). Nevertheless, we still require a better understanding of the neurophysiological and transcriptional basis of these social effects to account for the complexities of individual behavioral phenotypes.

In this experimental study, we investigated whether queen presence signaling (*i*.*e*., QMP exposure) during different periods in the life of adult honey bees affects foraging behavior and gene expression at the age of foragers, which is the last phase in the life of a worker bee (5, 6). We used synthetic QMP to elucidate the long-term consequences of QMP presence or absence by treating newly emerged workers, nurses, and foragers (1-19 days of age). When bees reached the foraging age at about three weeks of age, we analyzed gene expression in central and peripheral nervous system tissues thought to be critical for regulating worker behavior. We focused on the mushroom bodies, antennal lobes, and antennae to capture the entire pathway of odor perception, from pheromone binding in the antennae to processing in the antennal lobes to the mushroom bodies, where learning and memory and multimodal sensory integration occur (49-51). We focused on the foragers for two reasons. First, foragers play a fundamental role for the nutritional health of a colony, but it remains unknown whether and how QMP affects the gene expression of forager-aged bees. Second, queen replacement leads to a gap in brood production, and requires bees of forager-age to rear the brood of the new queen. We might, thus, expect bees that experienced queenlessness at a young age to show strong changes in gene expression when reaching forager age. Finally, we tested whether the QMP treatment affected foraging behavior by observing our focal bees and quantifying foraging activity. We expected that the QMP treatment would alter transcriptional activities in foragers. Moreover, if QMP treatment has long-term consequences, we expected them to be most pronounced when treatment occurred early in life, as younger workers also responded more strongly to QMP than nurse/forager-aged bees (42).

## Materials and Methods

### Colony Set-up

Three *Apis mellifera* observation colonies were established from three regular sized colonies prior to the start of experiments from August to October 2019, each containing approximately 2000-3000 workers of mixed ages, from the Johannes Gutenberg University apiary in Mainz, Germany. Each observation colony was headed by a naturally mated unrelated queen and all observation colonies had three frames, brood, pollen, and honey reserves.

### Sample Collection

One colony at a time was studied. Frames with ready-to-emerge workers were collected from the respective source colony to maintain a similar genetic composition in observation colonies. These frames were stored in a climate cabinet at 34°C overnight. Newly emerged workers were carefully marked with randomly assigned cohort-specific colored number tags on their thorax and enamel paint on the abdomen. Marked bees were introduced to the respective observation colony and collected once the desired age for treatment was reached.

Brood frame collection was done in a staggered sequence to allow collecting of forager-aged bees (at 19-days), nurse-aged bees (at 7-days), and newly emerged workers (at 1-day of age) at the same time (Fig. S1). Mixed cohorts of ca. 150 bees consisting of forager-aged (n = 50) and nurse-aged (n = 50) workers, and newly emerged (n = 50) workers were evenly distributed and introduced to their designated cages (12 cm x 12 cm) with (QMP+; 1/10 amount of strip brand (the equivalent of 1 queen, Bee equipment, UK) or without artificial QMP (QMP-). They were kept in their respective cage for 2 days, with access to *ad libitum* queen candy. Queen candy was made by mixing powdered sugar and warm water until it was the consistency of putty. Afterwards, all cohorts were re-introduced to their observation colony (Fig. S1). We filmed the foraging activity of each cohort for 3 days once they reached 21 days of age, until 24 days of age, at which age they should typically be foragers. Foraging is the last task a worker bee performs before her death (6). After filming (see below), bees from the different cohorts were collected from observation colonies when they were 24 days old, using Eppendorf tubes and immediately preserved in liquid nitrogen. We also collected forager-age bees that remained in observation colonies for the duration of the experiments, *i*.*e*. workers that were never kept in a treatment cage but were returned to observation hives immediately after emergence and marking (henceforth called “control foragers”). All samples were stored in -80°C freezer until further analysis.

### Video Analysis

Each colony entrance was recorded for 3 days to quantify the foraging behavior of experimental bees at the age of 21 days. We filmed the entrance of the colonies for 8 hours/day, from 9 am to 5 pm and, subsequently, we analyzed foraging by noting the time of entry and departure of marked foragers. We logged behavioral observations using VLC Media Player and statistically analyzed the differences between colonies and treatment groups using generalized linear models (GLM) with the negative binomial distribution. We performed pairwise comparisons between age groups using the general linear hypothesis test (GLHT) with Bonferroni correction for multiple testing.

### *Brain dissection* and *RNA extraction*

The heads from n = 42 workers were cut from the body and fixed on melted dental wax in a pre-chilled petri dish over ice. The antennae were cut off and stored in 100 µL of Trizol™ (Invitrogen, USA). The mushroom bodies and antennal lobes were removed by making incisions through the antennal base, eyes, compound eye, and ocellus (52). The cuticles, glands, retina and tissue around the brain were removed and the exposed tissues of the head were submerged with cooled bee saline (154 mM NaCl, 2 mM NaH_2_PO_4_, 5.5 mM Na_2_HPO_4_, pH 7.2). Each dissection, one bee brain dissected into 3 tissues, was completed in less than 5 minutes to prevent degradation of RNA. Brain parts were stored in 100 µL of Trizol™ in -80 °C for later RNA extraction using RNAeasy Mini Extraction Kit™ (Qiagen, Germany) according to the manufacturers protocol. RNA was extracted from foragers treated immediately after emergence (newly emerged; n = 12), treated at nurse age (n = 12), treated at forager age (n = 12), and control foragers n = 6 (2 per colony) from a total of 3 colonies (4 samples from each colony consisting of 2 QMP+ and 2 QMP- in foragers that were treated immediately after emergence, at nurse age, at forager age and 2 samples from each colony consisting of 1 QMP+ and 1 QMP- in control foragers).

### Transcriptome Analysis

Aliquots of RNA from each sample were sent to Beijing Genomics Institute (BGI) for sequencing using BGISeq to get 100 base pair (bp) paired-end reads, obtaining ∼45 Mio clean paired reads per sample sequencing. Reverse transcription to cDNA was performed as part of the library preparation by BGI (Fig S2). Clean reads without adaptor sequences were provided by BGI and quality checked using *FastQC* v.0.11.8 (53). Clean reads were aligned using *HiSat2* v.2.1.0 (54) under default settings to the most recent honey bee genome HvA3.1 (55), with a mapping ratio of more than 95% per sample. To count how many aligned reads mapped to genes, we used *HtSeq* v.0.11.2 (56) to generate count tables under the default parameters. Count tables were generated separately for each sample and complied separately for each tissue (*i*.*e*. antenna, mushroom bodies, and antennal lobes), treatment (*i*.*e*. QMP+/-), and life-stage when experiencing the treatment (*i*.*e*. forager-, nurse-, and newly emerged-age).

### Gene Expression Analysis

Gene expression differences were analyzed between treatments (QMP+ vs QMP-) for each tissue (mushroom bodies, antennal lobes, and antenna) and life-stage (newly emerged-forager, nurse-forager, forager-forager, and control-forager) using the R package *DESeq2* v.1.24.0 (57). Before the analysis, an additional filtering step was added to ensure that only genes with counts of at least 9 reads in n-1 of the smallest sample size were used in the gene expression analysis. We analyzed gene expression separately for each tissue and age. We tested the effect of treatment with QMP on gene expression by using the likelihood ratio test (LRT) approach whereby a full model with treatment (QMP+/-) and colony-ID as fixed factors is compared with a reduced model containing only colony-ID, taking into consideration colony effects. Genes were considered differentially expressed if the false discovery rate (FDR) corrected p-value was < 0.05. All analysis was completed in R v.4.1.0. We performed permutations to test whether the overlap in differentially expressed genes between tissues differed from what could be expected to overlap by chance. We generated a gene universe that had the same total number of genes (n = 12, 320) which contained random gene IDs and zeros. We created a vector for number of permutations we wanted to generate (n = 1,000). We performed automatic iterations with the number of DEGs in each gene list to get overlap values with our randomly generated list. We plotted histograms of the distribution generated by the permutations against the overlap to see if what we found was within (expected by chance) or outside (not expected by chance) the normal distribution.

We performed both gene ontology (GO) and KEGG overrepresentation analysis on each differentially expressed gene list with the clusterProfiler package (58), using the honey bee annotation that can be retrieved with the R package AnnotationHub (59). To further investigate QMP treatment effects on specific genes, we focused on candidate genes involved in key individual and social behaviors and traits. We compiled lists of genes and molecular pathways associated with *foraging behavior* and *division of labor* in honey bees (52, 60-74). Furthermore, we searched for genes that play roles in *aging* (52, 75-78), *immunity* (79, 80), and *reproduction* (33, 34), *i*.*e*. processes known to be affected by queen signals (see Tables S1 for a complete list of the candidate genes). We cross referenced all of our DEG lists, including the overlapped DEGs, to the candidate gene list to find genes of interest from previous studies.

## Results

### Influence of QMP on gene expression

Transcriptome analysis across the three tissues showed that QMP treatment early in life produced the strongest response. Despite the longest time period between treatment and sampling, workers treated with QMP as newly emerged bees showed the most pronounced differences in gene expression: a total of 1,398 genes were differentially expressed in all tissues (Fig. 1). In contrast, nurse-aged or forager-aged bees exposed to our QMP treatments only altered the expression of 80 and 67 genes across tissues, respectively (Fig. 1). Tissue-specific analyses revealed strongest transcriptomic shifts in the mushroom bodies with 760 DEGs in foragers treated as newly emerged bees (Fig. 1C). The vast majority of differentially expressed genes were significantly upregulated in bees that experienced a lack of the queen signal during the first days of their life (Fig. 1, 68% in MBs, 85.7% in AL, 91.7% in AT.). Interestingly, the opposite pattern was found in bees treated at nurse-age (Fig. 1). Here, most differentially expressed genes were upregulated in QMP+ bees (Fig. 1) (94% in AL and 75.8% in AT).

**Figure 1:**
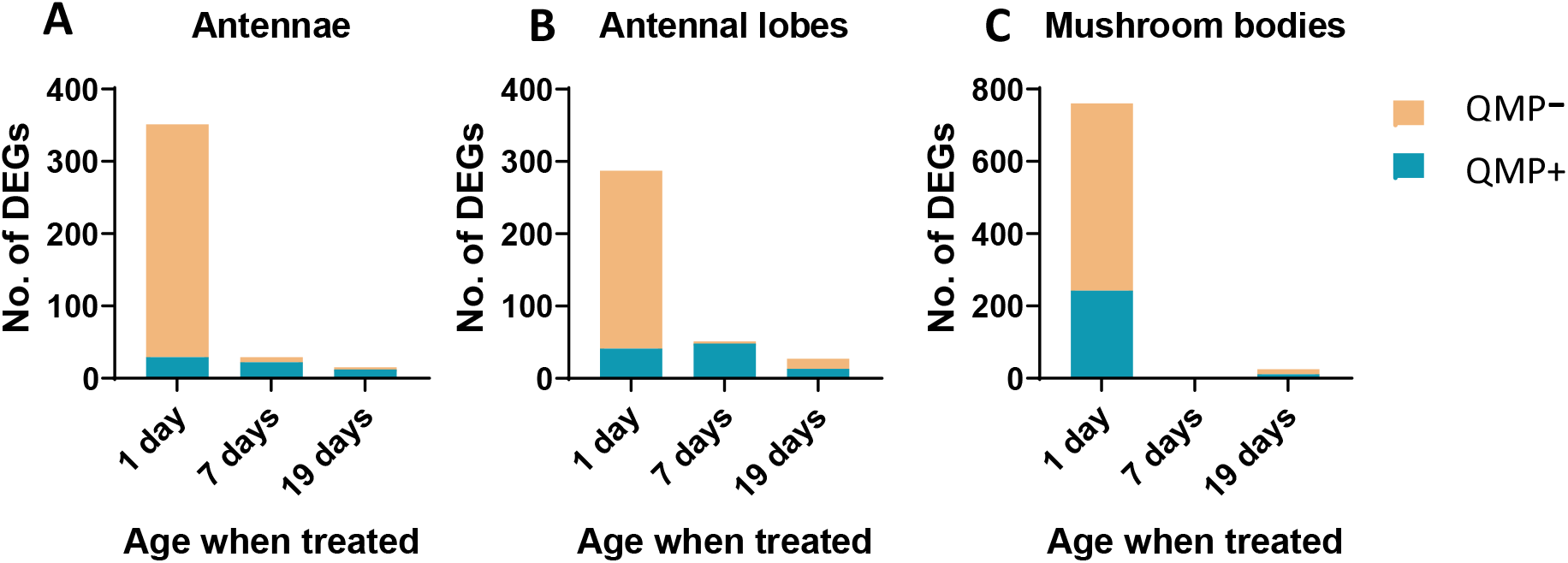
Number of differentially expressed genes: Differential gene expression across the different tissues (**A**) Antennae (**B**) Antennal lobes (**C**) Mushroom bodies. All bars indicate the total number of differentially expressed genes (DEGs) that are upregulated in each age group after QMP treatment (1 day = newly emerged, 7 days = nurse, 19 days = forager). Orange shading indicates samples treated 48h without exposure to QMP (QMP-) and blue shading indicates samples treated 48h with QMP exposure (QMP+).

We found the most enriched GO terms of the DEGs in foragers treated with QMP as newly emerged bees. Many of these GO terms were associated with RNA processing and binding in the antennal lobes and in the mushroom bodies (Table S2). We also found a number of behaviorally relevant candidate genes when comparing our DEGs with lists of genes found previously in the honey bee to be linked to foraging, division of labor, aging, reproduction and immunity genes (Table S1):

#### Mushroom bodies

Bees treated with QMP when newly emerged had differentially expressed genes associated with aging (*DNA repair proteins, telomere associated proteins*), division of labor (*kruppel homolog 1*), memory (*metabotropic glutamate receptor*), and foraging (*serotonin receptor, D2-like dopamine receptor*) (Table S1; Fig. 3). Bees treated with QMP at forager age had differentially expressed genes associated with cellular maintenance and DNA repair (Table S1).

#### Antennal lobes

Bees treated with QMP when newly emerged again had differentially expressed genes associated with aging (*telomere associated proteins* and *DNA repair proteins*) and learning (*Allatostatin A receptor*) (Table S1). Bees treated at nurse age had differentially expressed genes associated with foraging (e.g., *dopamine receptor, D1*) (Fig. 3). Bees treated at forager age had differentially expressed genes associated with learning (e.g., *Allatostatin A receptor*) (Table S1).

#### Antennae

Bees treated with QMP when newly emerged had differentially expressed genes associated with odor binding (*odorant receptor 4-like, general odorant-binding protein 71*) and aging (*DNA repair proteins*) (Table S1; Fig. 3). Bees treated with QMP at nurse age had differentially expressed genes associated with odor binding (*odorant receptor 53*) (Table S1; Fig.3). Bees treated with QMP as forager age had few (*15*) genes differentially expressed, many of which were uncharacterized.

#### Differentially expressed genes across tissues

Some genes were differentially expressed in more than one tissue. For the DEGs of the newly emerged bees, we found 15 genes that overlapped between the antennae and antennal lobes, 67 genes overlapped between the antennal lobes and mushroom bodies, 115 genes overlapped between the antennae and mushroom bodies, and 102 genes overlapped between all tissues (Fig. 2). Permutations showed that this gene expression overlap between all tissues, between mushroom bodies and antennae, and between mushroom bodies and antennal lobes were all outside of the range that could be expected by chance. However, the overlap between the antennae and antennal lobes was within the range of what could be expected by chance (Fig. S4). We found five genes that overlapped between the antennae and antennal lobes for samples treated with QMP at nurse-age. Note that there were no differentially expressed genes in the mushroom bodies. The gene expression overlap for nurse-age bees was within the range of what could be expected by chance. We found no overlap of DEGs between any tissues for samples treated at forager-age. All gene lists with a significant overlap were analyzed for matches against the candidate gene list (see Table S1) as well as GO enrichment analysis. We did not find any GO enrichments or matches against the candidate gene list.

**Figure 2:**
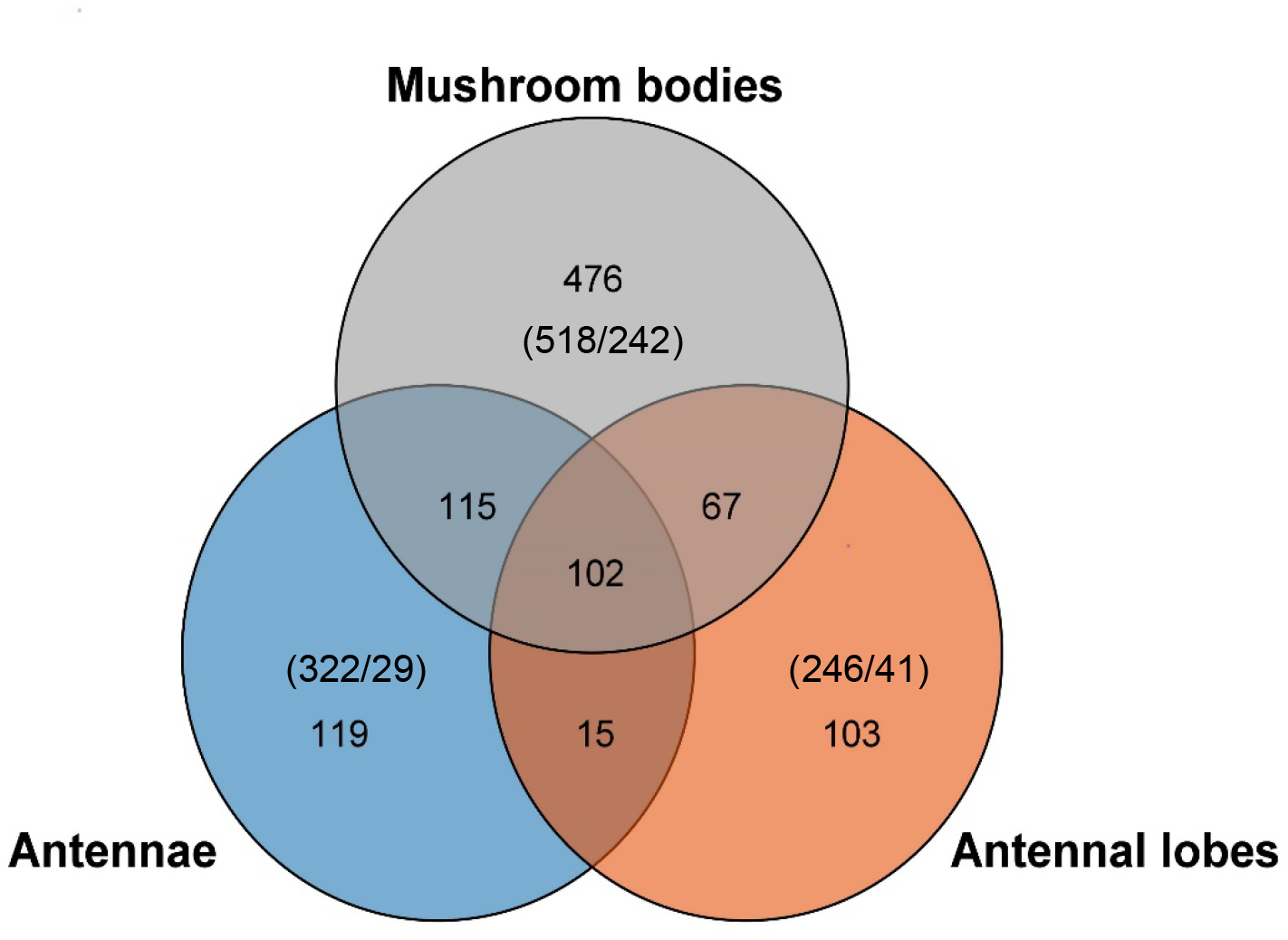
Gene overlap for workers with QMP treatment as 1 day newly emerged: Venn diagram representation of the overlap between DEGs of tissues for foragers treated with QMP treatment as 1 day newly emerged bees. Each color circle represents a different tissue (blue = antennae; orange = antennal lobes; grey = mushroom bodies). Numbers within each individual circle indicate the number of unique of DEGs for that specific tissue (i.e. 119 in the antennae; 103 antennal lobes; and 476 mushroom bodies). Numbers between circles indicate the number of shared DEGs between tissues (i.e. 15 between the antennae and antennal lobes; 67 between the antennal lobes and mushroom bodies; 115 between the antennae and mushroom bodies; 102 between all tissues). Numbers in parenthesis indicate the number of upregulated genes (QMP-/QMP+) in treated groups.

### Foraging behavior

We found an overall effect of age when treated with QMP on the sum of foraging trips (GLM, negative binomial distribution, Χ^2^ = 17.91, p < 0.001, Fig. 4) and average time spent foraging (LME, normal distribution, Χ^2^ = 12.25, p = 0.002, Fig. 4), but no effect of the QMP treatment (X^2^ = 0.259, p = 0.6; X^2^ = 0.705, p = 0.4 respectively) and no interaction between QMP treatment and age when treated (X^2^ = 2.603, p = 0.272; Χ^2^ = 4.18, p = 0.123 respectively, Fig. 4).

**Figure 3:**
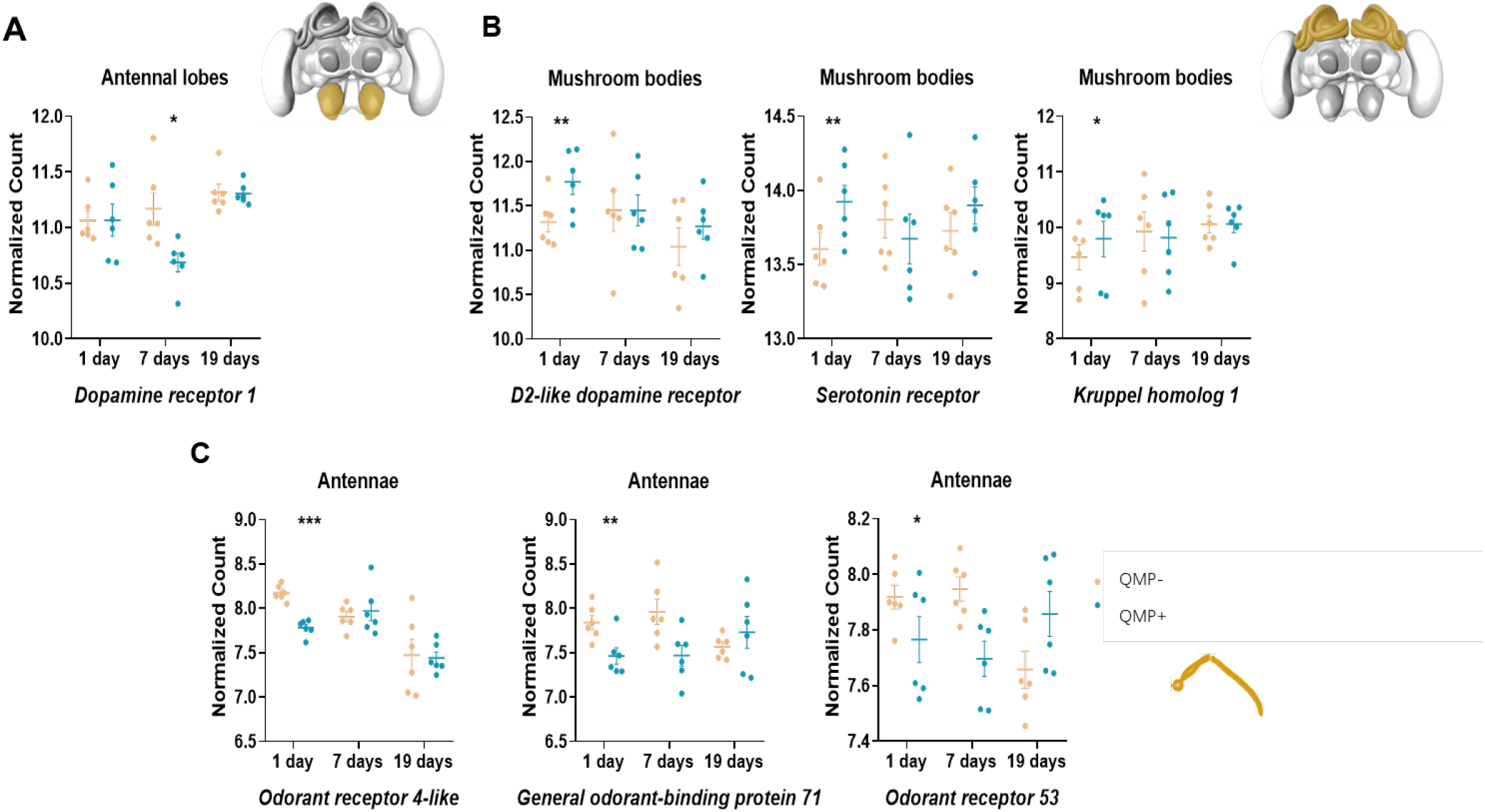
Genes of interest: Significant genes expression patterns for genes of interest. We searched the top differentially expressed genes (DEGs) of each gene list separately for genes associated with foraging and other behaviors (see Table S1). We show only the tissues where the gene was significant: In the (**A**) antennal lobes, only dopamine receptor 1 (Dop1) was significant (p = 0.044) in workers treated with QMP at 7 days nurses. In the (**B**) mushroom bodies, D2-like dopamine receptor (Dop3) (p = 0.004), Serotonin receptor (5-ht1) (p = 0.003), Kruppel homolog 1 (Kr-h1) (p = 0.038) in workers treated with QMP at 1 day as newly emerged bees. In the (**C**) antennae, odorant receptor 4-like (LOC107966034) (p < 0.001) and general odor binding protein 71 (LOC113218767) (p = 0.001) in workers treated with QMP at 1 day as newly emerged bees. Odorant receptor 53 (Or53) (p = 0.043) in workers treated with QMP at 7 days as nurses. All p-values shown are after FDR correction.

**Figure 4:**
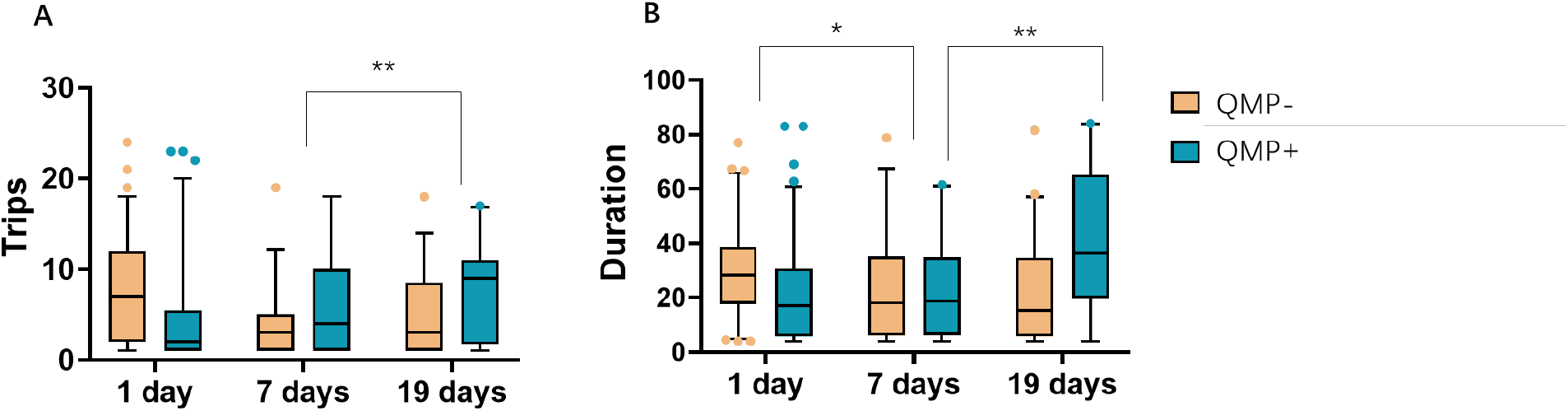
Effect of QMP exposure on foraging activity: Effect of QMP treatment on foraging activity (**A**) total number of foraging trips and (**B**) average foraging duration. On the y-axis, each age group when treated with QMP is represented (1 day = newly emerged; 7 days = nurse; 19 days = forager). Box and whisker plots are color coded according to treatment (orange = 48h without QMP treatment (QMP-); blue = 48h with QMP treatment (QMP+)). There was no difference in foraging activity between QMP treatment groups. Pairwise tests between age groups were performed using General Linear Hypothesis Tests (GLHT) and differences were found in the total number of foraging trips (7 days nurse vs. 19 days foragers GLHT: z-score - 3.214, p-value 0.00393.) and average foraging duration (1 day newly emerged vs. 7 days nurses GLHT: z-score -2.813, p-value 0.01340; and 7 days nurse vs. 19 days forager GLHT: z-score - 3.202, p-value 0.00385).

We performed pairwise comparisons of the age when treated with QMP for both total number of foraging trips and average foraging duration using Bonferroni correction for multiple testing and found that bees treated as nurses had fewer foraging trips and spent on average less time foraging than those treated at foraging age (General Linear Hypothesis Test (GLHT): z-value = -3.214, p = 0.003 and GLHT: z-value = -3.202, p =0.004 respectively; Fig. 4) and newly emerged bees (GLHT: z-value = -2.813, p = 0.013; Fig. 4), while there was no difference between newly emerged and nurse-age (GLHT: z-value = -2.120, p = 0.068; Fig. 4) in total number of foraging trips. Forager-age and newly emerged bees treated with QMP did not differ in total number of foraging trips (GLHT: z-value = -1.616, p = 0.106; Fig. 4) or average foraging duration (GLHT: z-value = -0.960, p = 0.6; Fig. 4).

## Discussion

Our results demonstrate a sensitive period of exposure to QMP in newly-emerged bees, which results in long-term transcriptional changes in the central and peripheral nervous systems. These molecular consequences persist across behavioral transitions until bees reach foraging age (24 days in this study). QMP elicits multiple distinct behavioral and physiological responses in workers, as both a releaser and primer pheromone, and thus produces responses on vastly different time scales (32, 81). The long-lasting effects of QMP exposure on gene expression in bees treated after emergence (1,398 genes in all tissues combined; Fig. 1) was most pronounced in the mushroom bodies (highest number of DEGs: 785) and, interestingly, most DEGs were upregulated when young bees experienced a lack of QMP exposure (Fig. 1). This suggests that a brief period of queenlessness early in life leads to extensive and persistent upregulation of transcription. In contrast, there were only small to moderate effects in bees treated later in life (80 genes in workers treated at 7 days of age (nurse age) and 67 genes in workers treated at 19 days (forager age); Fig. 1), suggesting that the time window for large-scale QMP effects closes very early in the life of a worker bee.

Sensitive periods exist across the animal kingdom and are generally characterized by a limited duration for key developmental processes in response to external cues or stimuli (44, 82, 83). An individual’s response to the cues or stimuli from the external environment is variable but the interaction can influence subsequent physiology and behavior. However, it is difficult to disentangle the role of the external environment in selecting for behavior in regards to learning plasticity versus the physiological traits of the individual in understanding how an organism responds to cues or stimuli. In humans, it is well-established that challenging conditions in utero, where an individual is buffered from the external environment, and early childhood shape adult coping strategies, behavior, and health (84-86).

This study provides evidence that the social signal, *i*.*e*. QMP exposure at a critical life stage (1-day-old, newly-emerged bees) is responsible for the previously observed worker phenotypes in response to pheromone signals. Young workers are more attracted to QMP, which results in the retinue response (18, 81, 87), while older workers engaging in foraging are repelled by queen pheromones (41, 88). This is consistent with the patterns in gene expression we found, where the transcriptional activity of 1-day newly emerged workers is drastically altered, while the transcriptional activity of 19-day foragers remains largely unchanged by QMP treatment (Fig. 1). Thus, forager-aged bees are largely resistant to QMP effects both in their behavior and upstream gene expression. Future studies could explore if long-term transcriptional effects observed during early life exposure to QMP are similar to pheromonal experiences during the larval and pupal stages.

QMP in high doses is reported to be repellent to bees (89, 90) and can make workers aggressive (91). QMP exposure during early adult life regulates the expression of D1-like, D2-like, and octopamine receptor genes (41, 42) which has been implicated in aversive learning in insects. We found that QMP exposure in early life led to long-lasting changes in the expression of D2-like dopamine receptor (*AmDop3*) and D1-like dopamine receptor (*AmDop1*) in the mushroom bodies and antennal lobes, respectively (Fig. 3; Table S1). It has been suggested that QMP’s effects on aversive learning may serve to prevent young workers that attend the queen from forming an association between the queen and any unpleasant effects of her pheromone (23), which is consistent with the observation that young workers are more attracted to QMP and why the formation of aversive olfactory memories is prevented (23, 41).

Dopamine receptors have a crucial role in a broad range of behaviors, such as motor function, sensory processing, arousal, and reward signaling (92-94). *Amdop3* is widely expressed in the brain in both adults and pupae, with a unique pattern of expression compared to the other subtypes *Amdop1* and *Amdop2* (95). Homovanillyl alcohol (HVA), a major component of QMP, has been shown to reduce the concentration of brain dopamine levels in the centers associated with learning and memory (*i*.*e*., mushroom bodies). HVA selectively activates *Amdop3*, which blocks aversive learning in workers (96), possibly promoting the retinue response as seen in the upregulation of *Amdop3* in the mushroom bodies of bees treated after emergence. Since we also find an upregulation of *Amdop1* in the antennal lobes of bees experiencing the absence of QMP at nurse age, this could suggest an early trigger of non-nursing behaviors. We also show a serotonin receptor (*5-ht1*) to be upregulated in the mushroom bodies of bees treated with QMP as newly-emerged bees, suggesting its role in learning and memory. *5-ht1* injection prior to olfactory conditioning resulted in reduced memory storage and retrieval (97-99), while high *5-ht1* levels in the central nervous system were associated with visual information processing in honey bee foragers.

Changes in QMP exposure also alter the morphology and gene expression in key brain tissues involved in olfaction, learning, and memory: the antennal lobes and mushroom bodies, under short-term QMP exposure conditions (42, 43, 100-102). The transcription factor *Krüppel-homolog 1* (*kr-h1*) is linked to hormone mediated social organization in honey bees, bumble bees, and ants (74, 103, 104). We found that QMP treatment induced an upregulation of *kr-h1* in the mushroom bodies of foragers that experienced no QMP exposure as newly-emerged bees. Experiments on young workers (<1 week) have shown that QMP exposure activates nursing genes and represses foraging genes, suggesting that QMP delays the transition between behavioral states by acting on *kr-h1* in the mushroom bodies (15). Although we do not find behavioral evidence to suggest that foraging activity was suppressed in 1 day newly emerged bees treated with QMP, future studies could use the evidence we provide to further investigate nursing behaviors after exposure to queenlessness.

QMP has varied effects on brain transcriptional activity depending on the behavioral state of a worker (15, 105) and our study shows that the pheromone information provided by QMP is processed differently at different adult stages. Accordingly, we find an upregulation in the expression of odor binding proteins (*odorant receptor 4-like* and *general odorant-binding protein 71*) and receptors (*odorant receptor 53*) in the antennae of workers treated with QMP as newly-emerged and nurse-aged bees (Fig. 3). A variety of odorant binding proteins (OBPs, ca. 21) and odorant receptors (ORs, ca. 180) have been characterized in the honey bee (60, 106) but their role in odor discrimination is still poorly understood. While we present further evidence for the role of some OBPs and ORs in the context of QMP signaling, more research is needed to understand their functioning in the context of odor differentiation and subsequent behavioral regulation.

In summary, we show an early sensitive phase for QMP exposure and long-lasting effects on transcriptional activity of the brain and antennae of young honey bee workers. We build on current knowledge that queen pheromones not only attract young workers and affect gene expression, but that these changes can persist into late foraging age. We observed the strongest changes in the mushroom bodies in workers that experienced an absence of QMP in early life. The results suggest an impact on learning and memory, but surprisingly not on foraging activity. Future research should observe whether these long-lasting changes in gene expression triggered by early life treatment alter behaviors such as nursing or immune function as suggested by the observed gene expression patterns presented here and in previous studies (Fig. S5).

## Supporting information

Candidate gene BLAST results

GO enrichment results

## Acknowledgements

We would like to thank the Deutsche Forschungsgemeinschaft that funded this project with a grant to C. Grüter and S. Foitzik (GR 4986/3-1 and FO 298/27-1). We are grateful to Romain Libbrecht, Marah Stoldt and Francisca Segers for their support in the gene expression analyses. Our laboratory work was supported by Marion Kever and Jenny Fuchs.

## Author Contributions

A.K., T.P., S.F. and C.G. conceived the study and designed the experiments. T.P. conducted the field experiments, honey bee brain dissections, RNA extraction, and gene expression analysis. Data was supported by S.F and C.G. All authors contributed to writing the manuscript.

## Data Accessibility

The data that support the findings of this study are available from the corresponding author upon reasonable request. All sequencing data is deposited in the BioProject database under the BioProject ID PRJNA792844.

## Supplemental Tables and Figures

**Figure S1:**
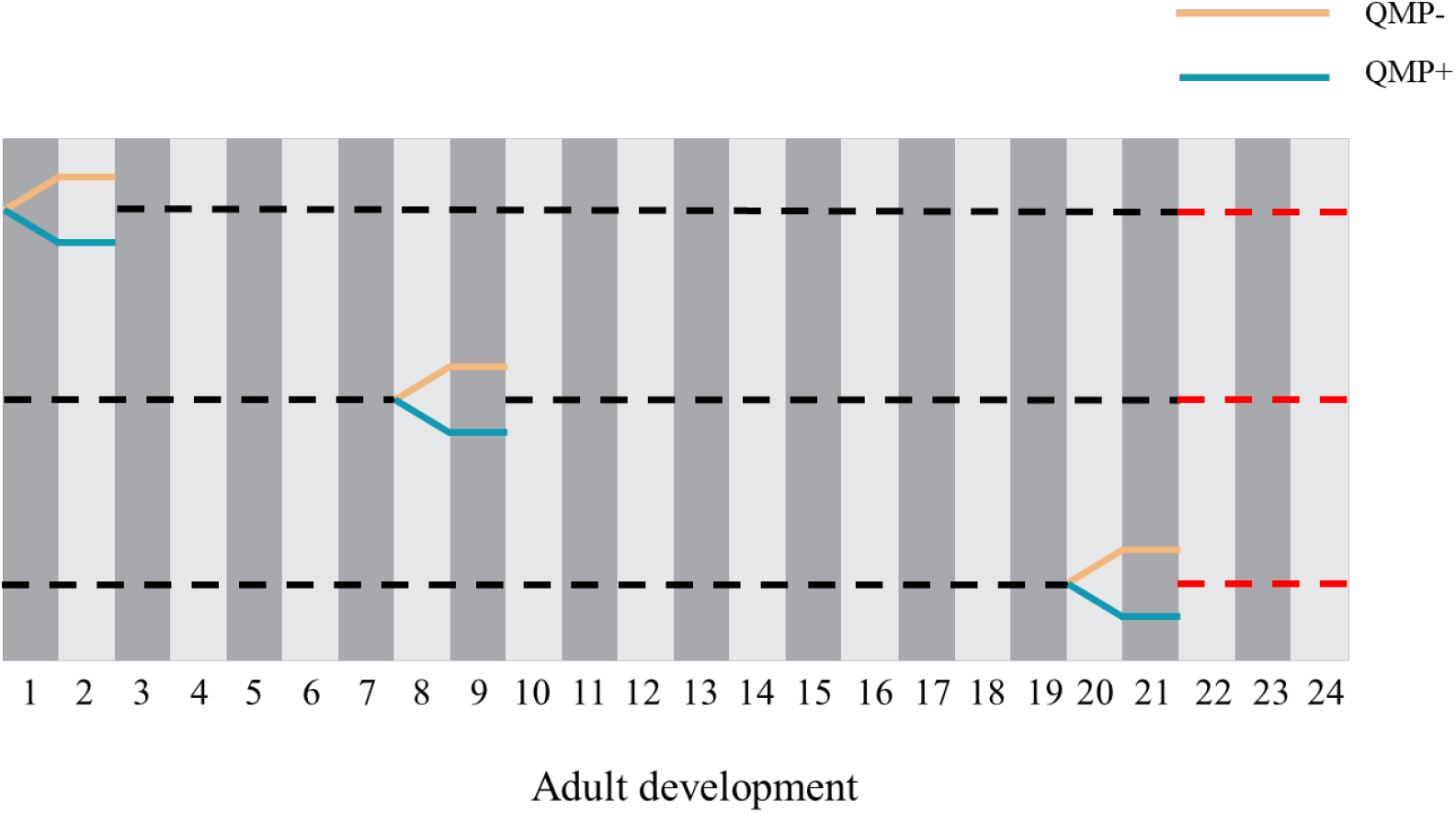
Experimental design: Experimental design overview. The black dotted line represents when workers were in the observation colony; the solid line represents when the workers were in the cage for treatment; the red dotted line represents when the workers were filmed in the observation colony.

**Figure S2:**
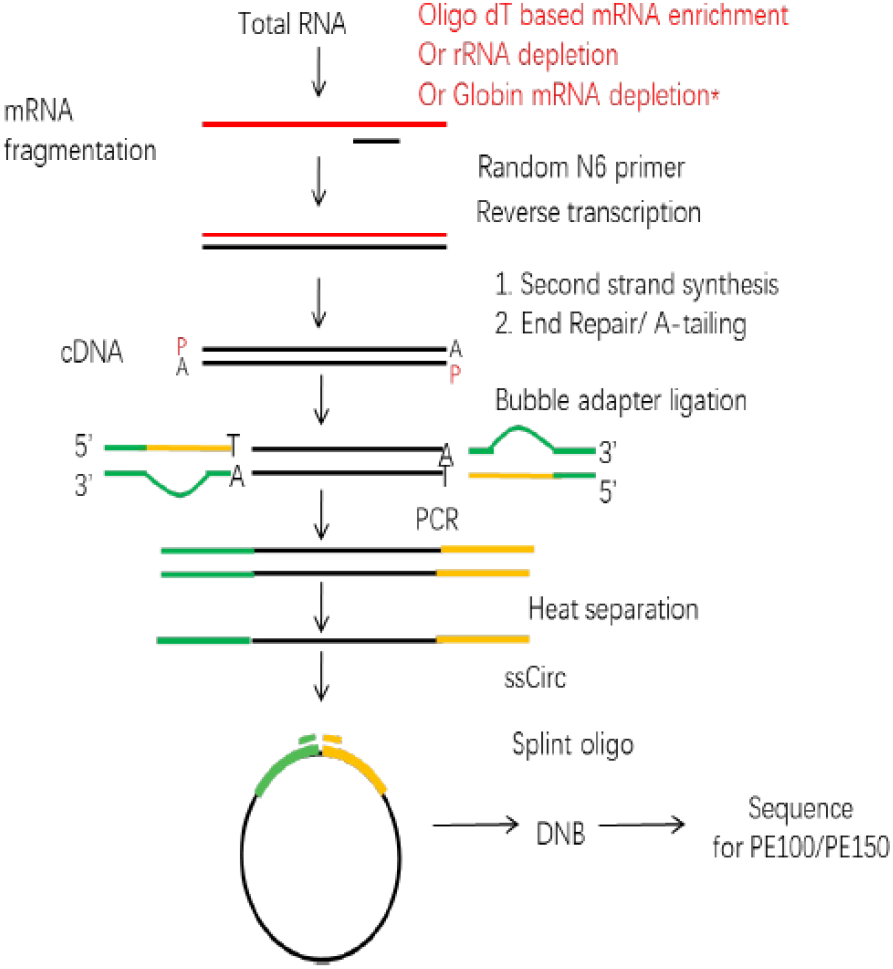
RNAseq pipeline: RNA sequencing workflow provided by Beijing Genomic Institute (BGI). Total RNA was extracted from 5 tissues (antennae; antennal lobes; mushroom bodies; central brain; and subesophageal ganglion) and sent to BGI for sequencing using BGISeq to get 100 base pair (bp) paired-end reads. We obtained ∼45 Mio clean paired reads per sample sequencing.

**Figure S3:**
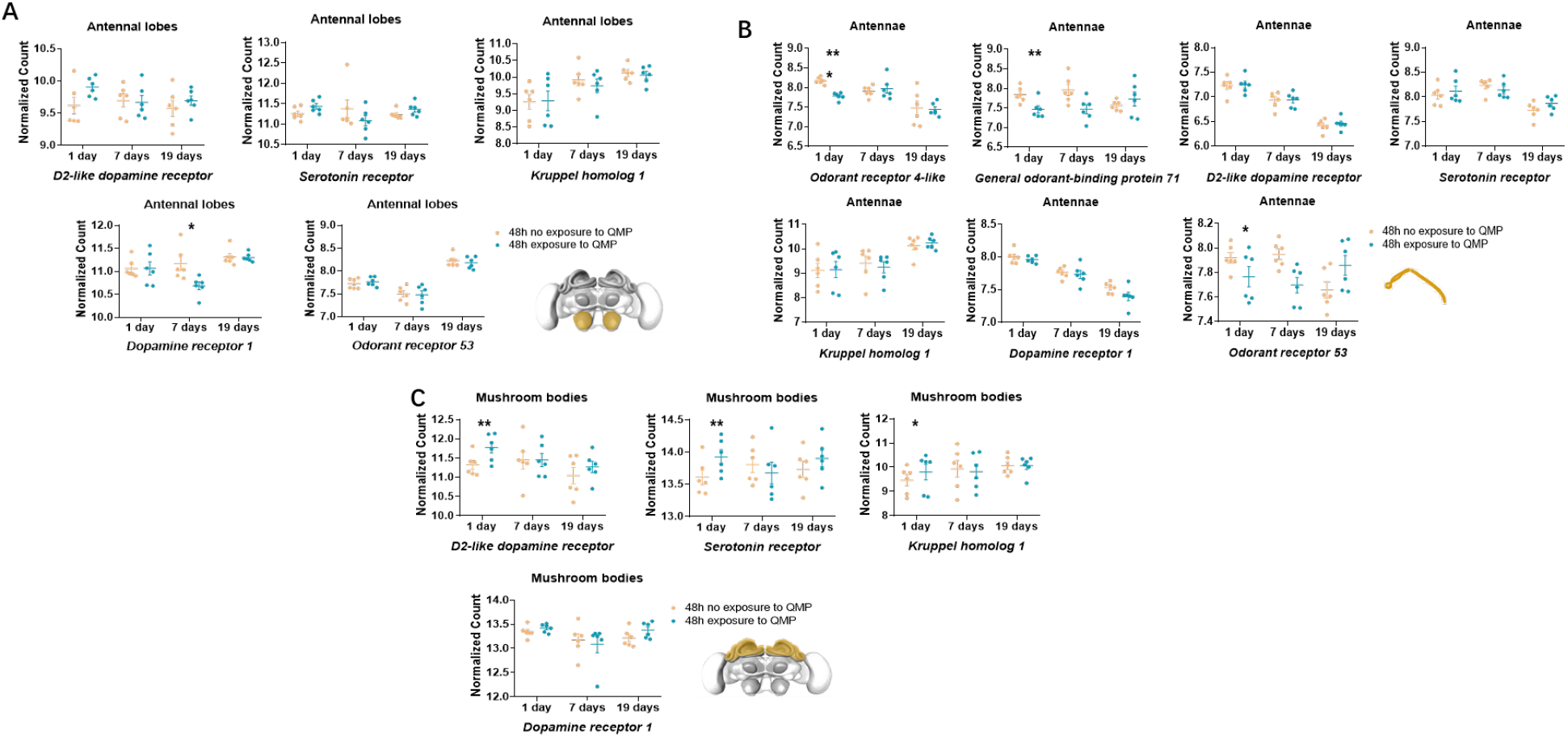
Normalized read counts for genes of interest for all tissues: Gene expression patterns for genes of interest for all tissues (**A**) antennal lobes; (**B**) antennae; and (**C**) mushroom bodies. We show the gene expression patterns from Fig. 3 as well as the non-significant tissues.

**Figure S4:**
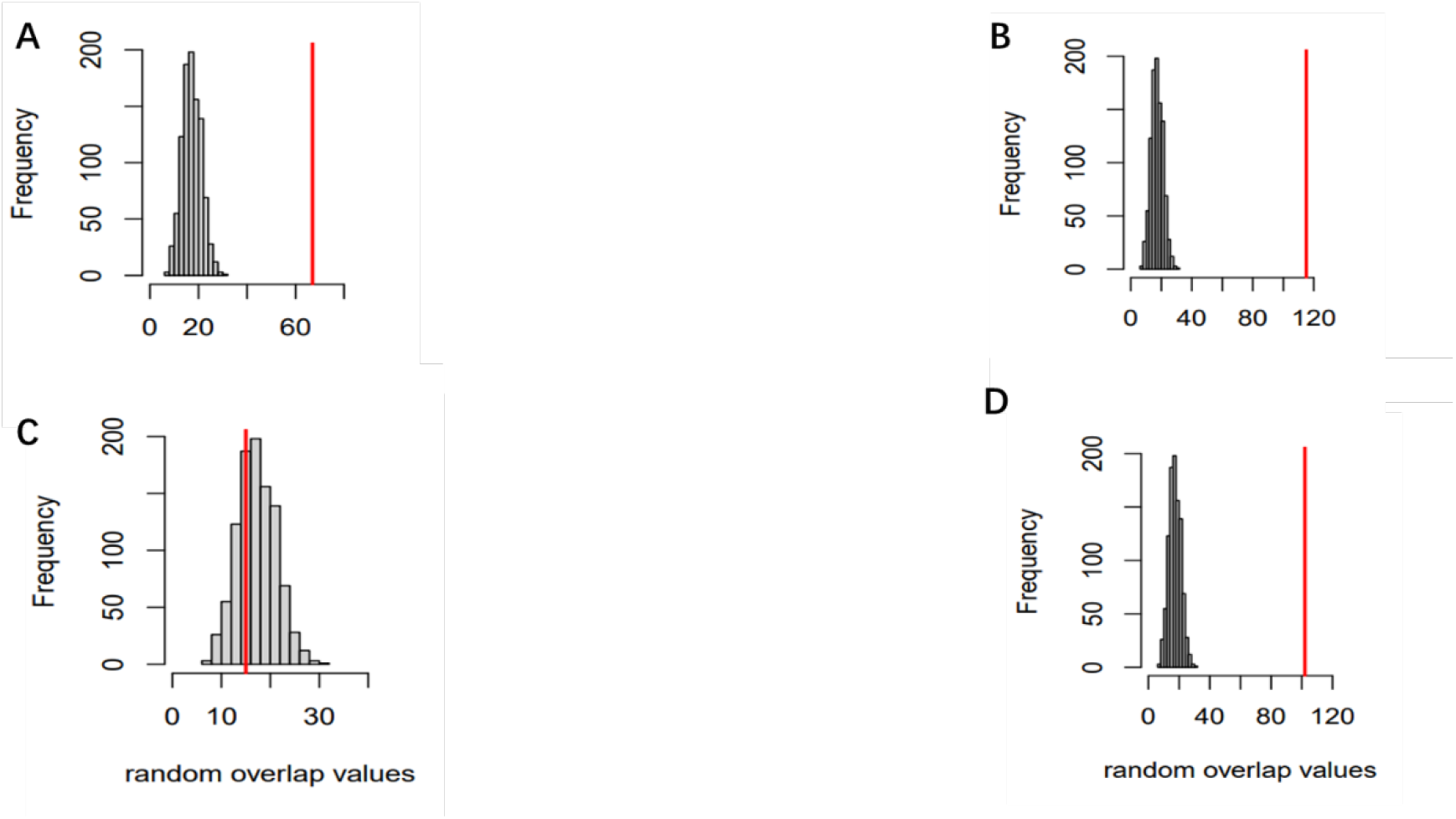
Gene expression overlap results: Permutation results for the gene expression overlap between tissues for samples treated with QMP as 1 day newly emerged bees. (**A**) mushroom bodies vs. antennal lobes (n = 67); (**B**) mushroom bodies vs. antennae (n = 115); (**C**) antennae vs. antennal lobes (n = 15); and (D) all tissue overlaps (n = 102). Red bar indicates the number of overlapped genes we found.

**Figure S5:**
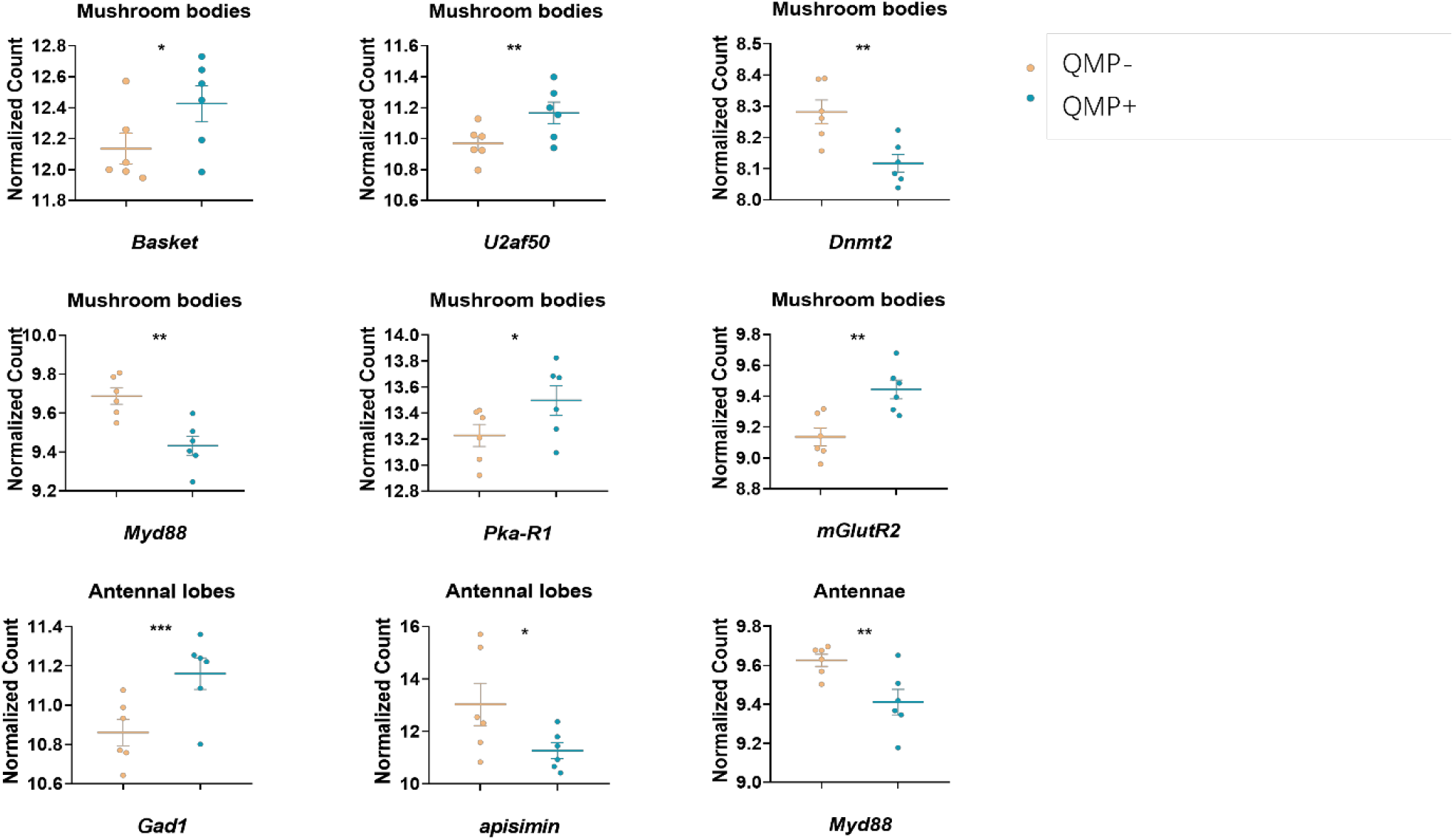
Candidate genes of interest: Genes of interest from the candidate gene blast search. We compared all of our differentially expressed gene (DEG) lists to DEGs found in previous studies (see complete list table S1). Here we show only significant genes of interest in 1 day newly emerged workers treated with QMP. In order from top left: Basket (p = 0.047); U2af50 (p = 0.0082); Dnmt2 (p = 0.0043); Myd88 (p = 0.0033); Pka-R1 (p = 0.023); mGlutR2 (p = 0.0098); Gad1 (p = 0.00046); apisimin (p = 0.011); and Myd88 (p = 0.0072).

**Table S1: Candidate gene BLAST results:** we compiled a list of candidate genes associated with processes such as foraging behavior, division of labor, aging, immunity, and reproduction from the literature. We compared the candidate gene list to all of our differentially expressed gene (DEG) lists to find candidate genes of interest that matched previous studies.

**Table S2: GO enrichment results:** we performed a gene ontology (GO) enrichment analysis and KEGG overrepresentation analysis on each differentially expressed gene (DEG) list with the clusterProfiler package, using the honey bee annotation that can be retrieved with the R package AnnotationHub.

